# A Cellular Cytotoxicity Assay using Ready-to-Thaw Target Cells without Washing Steps

**DOI:** 10.64898/2026.01.21.700863

**Authors:** Jonathan Storm, Neele Kusch, Marina Güttler, Carla Fode, Lara Breuer, Jan Bartling, Cornelius Knabbe, Barbara Kaltschmidt, Christian Kaltschmidt

## Abstract

Functional testing of cytotoxic lymphocytes is essential for research and quality control (QC), but most assays require freshly prepared target cells and extensive handling. A ready-to-thaw, no-wash, flow cytometry–based cytotoxicity assay was developed using pre-labeled K562 targets cryopreserved in STEM-CELLBANKER® EX (SCB) as suitably sized aliquots. SCB tolerability was evaluated in K562, NK-92, and primary natural killer (NK) cells; post-cryopreservation label stability of CellTrace™ Violet (CTV) and carboxyfluorescein succinimidyl ester (CFSE) was assessed; freezing and thawing conditions were optimized; and wash versus no-wash workflows were compared using viability-based and absolute-count readouts, across effector-to-target (E:T) ratios with NK donors and NK-92 cells. Effector viability remained high at SCB concentrations up to 10%, and 5% SCB was selected for assay design. After cryopreservation, CTV labeling remained stable over the tested storage period, whereas CFSE showed substantial signal loss. Warm-medium thawing performed comparably to water-bath thawing, and the consolidated protocol (SCB plus fetal calf serum and thermal buffering) maintained high post-thaw target viability and recovery. In killing assays, lysis increased with increasing E:T ratios; omission of the post-thaw wash had minimal impact, and 5% SCB did not impair cytotoxic function. This ready-to-thaw workflow reduces hands-on time and sample manipulation, while improving standardization for reproducible results and enabling high-throughput functional testing and QC.

## 1. Introduction

Cytotoxic lymphocytes, including T cells and natural killer (NK) cells, have emerged as highly promising candidates for the immunotherapy of cancer and viral infections [1–7]. The use of antigen-specific T cells, particularly in the form of chimeric antigen receptor (CAR)-T therapies, has revolutionized the treatment of hematological malignancies, and several are already available as approved therapies [8–10]. In parallel, NK cells provide antigen-independent cytotoxicity, offering an attractive “off-the-shelf” platform that may be further enhanced by CAR engineering [11,12]. Both modalities critically depend on rigorous assessment of their cytolytic function, not only during preclinical development but also as part of manufacturing and quality control (QC) pipelines for clinical-grade cell therapies [13– 15].

A variety of assays are available to evaluate lymphocyte-mediated cytotoxicity in vitro. The chromium-51 (51Cr) release assay, described in 1968 [16], has long been considered the gold standard for measuring NK- and T cell cytotoxicity [17,18]. However, concerns related to radioactive waste, safety, and disposal have motivated the development of non-radioactive alternatives, such as lactate dehydrogenase (LDH) release, luciferase-based readouts [19], or fluorescence-based assays evaluated via microscopy, photometry, or flow cytometry [20]. Analysis of functional parameters like the expression of activating and inhibitory receptors [21], degranulation (e.g. through CD107a/LAMP1 surface exposure) [22,23], and the production of interferon-γ [24] are widely used to assess activation status, but they do not directly report target cell death. While calcein-AM release assays are widely applied, they suffer from variable dye loading efficiencies, high spontaneous leakage, and incomplete release from apoptotic bodies, which may lead to the underestimation of true target cell lysis [17]. In contrast, fluorescent dyes such as CellTrace™ Violet and carboxyfluorescein succinimidyl ester (CFSE) can be used to label target cells [17,25] and are often used in combination with DNA-binding viability dyes like propidium iodide (PI) to determine dead cells in flow cytometry–based assays [26]. This approach enables direct discrimination of effector and target cells, robust assessment of target cell viability, and quantification of specific lysis at the single-cell level. Although flow cytometry–based assays are comparatively time-consuming, their sensitivity, versatility, and compatibility with multiparametric analysis make them particularly suitable for both research and translational applications.

Despite these methodological advances, a persistent limitation remains: cytotoxicity assays require fresh preparation of target cells. This may include harvesting, labeling, and quality control, all on the day of the experiment. This not only creates a high workload, but also complex, lengthy protocols, in consequence making operator fatigue a probable source of error and also reducing reproducibility across experiments. A genuine ready-to-thaw killing assay format—including pre-labeled, cryo-preserved target cells that can be used directly without additional processing steps like washing—is currently unavailable. Commercially, only partial solutions exist: kits that provide dyes for user-based staining [27–29], or genetically engineered reporter cell lines that express tags for luciferase based detection [30]. While these approaches are useful, they do not offer standardized, pre-stained, small-size aliquoted and cryopreserved target cells for flow cytometry–based cytotoxicity readouts. Thus, a gap remains for a practical and reproducible assay format that could streamline functional testing in immunotherapy research and QC.

Recent advances in cryopreservation have produced media formulated from infusion-grade, intravenously used inactive ingredients. STEM-CELLBANKER^®^ EX (SCB) is marketed for direct-use workflows (no pre-infusion washing) and is compatible with simple freezing procedures, making it suitable to be present at low residual levels during biological processes [31,32]. Importantly, the cryopreservation medium must be non-toxic at low concentrations, and minimal irritation is essential to preserve immune-cell function in its presence. These properties enable the development of pre-labeled, cryo-preserved, non-washed target cells for flow-cytometric cytotoxicity assays.

To operationalize this concept in standard flow-cytometry workflows, several practical constraints informed the assay design. Non-permeant DNA dyes (e.g., PI/DAPI) are compatible with no-wash protocols but remain susceptible to time-dependent uptake during acquisition, whereas fixable viability dyes require washing because effective quenching of unbound dye is unavailable. Acquisition is typically limited by flow rate (<100 µL/min, often ∼60 µL/min), so low per-µL cell concentrations prolong measurement and can bias live-cell readouts. To achieve a 96-well–compatible format while retaining compatibility with standard flow-cytometry hardware, we chose 1.1 mL Micronic tubes. With an upper working volume of ∼1 mL per tube (to avoid transfers) and downstream additions (e.g., EDTA and a viability dye), the co-culture volume was constrained to ∼800 µL. Accordingly, the dilution required to achieve low residual cryomedium set the maximum feasible frozen aliquot volume. To ensure unambiguous enumeration, targets (not effectors) were labeled, and a brief EDTA step was used to disrupt conjugates prior to acquisition [25]. These constraints define the design space evaluated in the present study.

In this study, we demonstrated the feasibility of a “no-wash, ready-to-thaw flow cytometry–based cytotoxicity assay”. Using NK-92 cells and primary NK cells as models of antigen-independent effector function, we systematically evaluated (i) the tolerance of effector and target cells to SCB, (ii) the feasibility of omitting washing steps after thawing, (iii) the stability of fluorescent target cell labeling after cryopreservation, and (iv) workflow optimizations such as small-volume aliquoting. Together, these advances provide the proof of principle for a streamlined, reproducible, and translationally relevant assay format that addresses a critical gap in cytotoxicity testing.

## 2. Materials and Methods

### 2.1 Cell Lines and Primary Cells

K562 cells (ACC-10, DSMZ–German Collection of Microorganisms and Cell Cultures, Leibniz Institute, Braunschweig, Germany) were cultured in RPMI-1640 medium (Sigma-Aldrich, Schnelldorf, Germany, or PAN-Biotech, Aidenbach, Germany) supplemented with 10% fetal bovine serum (FBS; VWR, Darmstadt, Germany, or PAN-Biotech), 2 mM L-glutamine (Sigma-Aldrich), and 100 U/mL penicillin plus 100 µg/mL streptomycin (Sigma-Aldrich). Cultures were maintained by medium supplementation every 2–3 days and passaged once or twice weekly.

NK-92 cells (ATCC^®^ CRL-2407™, American Type Culture Collection, Manassas, VA, USA) were cultured in MEM-α (Sigma-Aldrich) with 12.5% fetal calf serum (FCS; PAN-Biotech), 12.5% horse serum (PAN-Biotech), 100 U/mL penicillin plus 100 µg/mL streptomycin (Sigma-Aldrich), and 100 U/mL recombinant human Interleukin-2 (IL-2) (Proleukin S, Novartis, Basel, Switzerland). Cultures were maintained as above; seeding density was 0.2–1 × 10^6 cells/mL.

Primary NK cells were isolated from buffy coats of healthy donors (Institut für Laboratorium-sund Transfusionsmedizin, HDZ NRW, Bad Oeynhausen, Germany), processed in accordance with the German Hemotherapy Guidelines. NK-cell enrichment used RosetteSep™ Human NK Cell Enrichment Cocktail with SepMate™-50 tubes and Lymphoprep™ (all Stemcell Technologies, Cologne, Germany), per manufacturer’s instructions. Residual erythrocytes were lysed with EasySep™ Red Blood Cell Lysis Buffer (Stemcell Technologies). Purified cells were cultured in NK-MACS^®^ medium (Miltenyi Biotec, Bergisch Gladbach, Germany) with 5% human AB serum (PAN-Biotech), NK-MACS^®^ supplement (Miltenyi Biotec), 100 U/mL penicillin plus 100 µg/mL streptomycin (Sigma-Aldrich), and 500 U/mL recombinant human IL-2 IS (Miltenyi Biotec). For SCB toxicity analysis, NK cells were seeded at 1 × 10^6 cells/mL and cultured for two weeks (0–100%) or overnight (0–25%) in standard tissue culture flasks (Sarstedt, Nümbrecht, Germany). For cytotoxicity assays, NK cells were expanded in G-Rex^®^ 24-Well Plates (Wilson Wolf, Saint Paul, MN, USA) for 60–62 days as described previously [33].

### 2.2 Flow Cytometric Cell Counting

Absolute concentrations were determined using CountBright™ absolute counting beads (Thermo Fisher Scientific, Darmstadt, Germany) on a Gallios flow cytometer (Beckman Coulter, Krefeld, Germany) or by the integrated volumetric function of a Northern Lights flow cytometer (Cytek, Fremont, CA, USA). For viability, cells were stained with PI (1:2000 to 1:1000; BioLegend, Koblenz, Germany) or 4’,6-diamidino-2-phenylindole (DAPI; 1:10,000; BioLegend). Staining was performed in PBS (PAN-Biotech) with 2 mM EDTA (Carl Roth, Karlsruhe, Germany) and 2% FBS (PAN-Biotech), hereafter PBSE2%FCS. PI was used either alone or with CellTrace™ Violet (CTV), CellTrace™ Far Red (CTFR), or CellTrace™ CFSE (all Invitrogen, Thermo Fisher Scientific, Darmstadt, Germany), and in CFSE-based assays when samples were acquired on Northern Lights. In CFSE assays on the Gallios, DAPI was used instead of PI. Where CTV/CTFR/CFSE were used, only dye-positive targets were counted. On the Gallios, 50 µL CountBright™ beads were added per sample. Data were analyzed in Kaluza (Beckman Coulter) or SpectroFlo (Cytek). Calculation of absolute counts and specific lysis is detailed in Section 2.15. Instrument configuration and gating are in Section 2.14.

### 2.3 K562 Labeling

For CTV, CFSE, and CTFR labeling, cells were resuspended at 1 × 10^6 cells/mL in PBS and incubated with dye (1:2000 for CTV/CTFR; 1:8000 for CFSE, or 1:4000 for CFSE in cytotoxicity assays; all Invitrogen, Thermo Fisher Scientific) for 15 min at 37 °C in the dark. Then 10 mL pre-warmed medium were added, incubated 5 min at 37 °C, centrifuged (300 × g, 5 min), washed twice, and resuspended in complete medium or cryopreservation solution.

### 2.4 Toxicity of STEM-CELLBANKER^®^ EX During 24 h Culture

To evaluate SCB cytotoxicity, K562, NK-92, or primary NK cells (cultured for two weeks [0–100%] or overnight [0–25%]) were incubated for 24 h in culture medium supplemented with SCB (Amsbio, Frankfurt am Main, Germany). K562 used Roswell Park Memorial Institute 1640 (RPMI-1640), NK-92 used Minimum Essential Medium, alpha modification (MEM-α), and primary NK used NK-MACS^®^ medium (as in Section 2.1). Cell counts were assessed by flow cytometry (Section 2.2). Cells were seeded with medium + SCB to final concentrations of 0–25% or 0–100% (v/v) in small-volume tubes (Sysmex, Görlitz, Germany) at 50,000 cells/100 µL. After overnight culture, cells were stained with 100 µL PI solution (2:1000 in Phosphate-buffered saline (PBS) with 2mM Ethylenediaminetetraacetic acid (EDTA) and (PBSE)2%FCS) plus 50 µL CountBright™ beads for absolute counting (Sections 2.2, 2.15).

### 2.5 Evaluation of Density Effects and Washing in Cryopreserved K562 Target Cells

K562 cells were harvested, counted by flow cytometry (Section 2.2), pelleted, and adjusted to 1–5 × 10^6 cells/mL in SCB. Aliquots (1 mL) were transferred to CryoPure tubes (Sarstedt) and placed in a styrofoam box at −80 °C for 24 h, then moved to liquid nitrogen (−196 °C). For evaluation, vials were rapidly thawed at 37 °C to a small ice crystal. Half of the suspension was used without washing; the other half was washed (dilution with warm medium, 300 × g, 5 min) and resuspended to the original volume in K562 medium. Cell counting followed Section 2.2.

### 2.6 CTV Fluorescence Stability Over Time

Cells were stained with CTV (Section 2.3), cryopreserved/thawed (Section 2.5), and analyzed by flow cytometry (Section 2.2) on days 6, 8, and 13 after cryopreservation.

### 2.7 Direct Comparison of Washed and No-Wash Flow Cytometric Cytotoxicity Assays

Cryopreserved targets were thawed (Section 2.5) and split into no-wash and washed fractions (washed: dilution with warm medium, 300 × g, 5 min, resuspension to equal volume). Fractions were adjusted to identical target densities and co-incubated with NK-92 or primary NK cells at E:T = 0:1, 1:1, 3:1, 9:1 in Sysmex low-volume tubes. After 4 h at 37 °C, 5% CO_2_, samples were cooled to 8 °C for 30 min, supplemented with 2 mM EDTA and PI (1:1000 in PBSE2%FCS), and 50 µL CountBright™ beads. Acquisition: Gallios; analysis: Kaluza. Specific lysis was calculated as in Section 2.15.

### 2.8 Comparison of Different Small-Volume Thawing Conditions

K562 cells (5 × 10^6 cells/mL in SCB) were aliquoted at 10, 20, 40, 80 µL (Micronic tubes) and frozen at −80 °C in a styrofoam box. Thawing used direct addition of 37 °C medium at 1:4 (aliquot:medium) (final 20% SCB) or rapid water-bath thaw to a small ice crystal followed by dilution. Cells were gently resuspended and analyzed for viability on the Gallios (Section 2.2).

### 2.9 Assessment of Post-Thaw Viability Across Freezing Volumes (Direct-Thaw Method)

K562 cells (5 × 10^6 cells/mL in SCB) were aliquoted at 10, 20, 40, 80, 120, 160 µL (Micronic) and frozen at −80 °C in a styrofoam box. Thawing was by direct addition of 37 °C medium at 1:4. Cells (labeled with CTV, CTFR, or CFSE) were resuspended and analyzed for viability on the Gallios (Section 2.2).

### 2.10 Assessment of FCS Supplementation and Automated Dispensing (MANTIS^®^)

K562 cells (1 × 10^6 cells/mL in PBS) were stained with CFSE (Section 2.3), counted (Section 2.2), and resuspended in SCB at 5 × 10^6 cells/mL, either without additives or +20% FCS. Suspensions were dispensed manually or via MANTIS^®^ Liquid Handler (Formulatrix^®^). Aliquots of 40 µL were placed into Micronic tubes in a 96-well rack and cryopreserved/thawed as in Section 2.8. Post-thaw viability and CFSE mean fluorescence intensity (MFI) were measured on Northern Lights using PI (Section 2.2).

### 2.11 Evaluation of Isopropanol for Thermal Buffering During Cryopreservation

K562 cells were harvested, stained, and resuspended in SCB + 20% FCS as in Sections 2.3, 2.8. Aliquots of 40 µL (Micronic) in a 96-well rack were frozen either directly in a styrofoam box at −80 °C or surrounded by heat-sealed, isopropanol-filled plastic bags within the box. Thawing followed Section 2.8. Post-thaw viability/recovery were analyzed on Northern Lights (Section 2.2).

### 2.12 Validation of Optimized Small-Volume Cryopreservation

K562 cells were stained with CFSE and resuspended in SCB + 20% FCS (Sections 2.3, 2.8). Aliquots of 40 µL were cryopreserved using isopropanol thermal buffering (Section 2.11). Thawing used direct addition of warm medium (Section 2.8). Post-thaw viability/recovery were analyzed on Northern Lights using PI (Section 2.2).

### 2.13 Cellular Cytotoxicity Assay Using Ready-to-Thaw Target Cells Without Washing

CFSE-stained K562 targets were cryopreserved in 40 µL aliquots as in Section 2.12, at 7.15 × 10^6 cells/mL (instead of 5 × 10^6 cells/mL). After thawing by direct addition of warm medium without washing (volumes per tube: 760 µL for 0:1, 720 µL for 1:1, 640 µL for 3:1, 400 µL for 9:1), NK-92 effector cells in K562 medium at 5 × 10^6 viable cells/mL were added directly (0 µL, 40 µL, 120 µL, 360 µL, respectively). Co-culture: 4 h at 37 °C, 5% CO_2_. Samples were cooled to 8 °C for 30 min, then supplemented with 50 µL PBSE2%FCS containing 34 mM EDTA and PI (6.8:1000; BioLegend). Acquisition: Northern Lights; absolute counts from the volumetric device. Specific lysis was calculated by viability-based and absolute methods (Section 2.15).

### 2.14 Flow Cytometer Configuration and Gating Strategy

Gallios channels/filters: CTV FL9 (405 nm; 480 DCSP–450 BP50), DAPI FL9 (405 nm; 480 DCSP–450 BP50), CFSE FL1 (488 nm; 550 DCSP–525 BP40), CTFR FL6 (638 nm; 710 DCSP–660 BP20), PI FL3 (488 nm; 655 DCSP–620 BP30). Compensation in Kaluza.

Northern Lights: 488 nm excitation, full spectrum; unmixing in SpectroFlo 3.1.0 (11142022) using single-stained controls and an autofluorescence reference.

Gating order: FSC-A vs Time → SSC-A vs FSC-A (cells) → FSC-H vs FSC-A (singlets) → fluorescence channels (target/effector, viability). On the Gallios, two free fluorescence channels were reserved for CountBright™ beads. Software: Kaluza for Gallios™ v1.2, Kaluza for Analysis v1.2; SpectroFlo v3.1.0.

### 2.15 Calculation of Specific Lysis and Absolute Counts

Control. The 0:1 condition (targets without effectors) after identical incubation served as control. Specific lysis (viability-based).

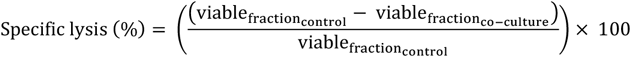

Specific lysis (absolute counts; beads or volumetric).

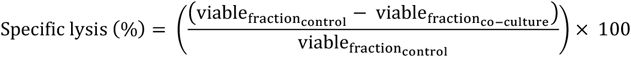

### 2.16 Data Processing and Statistics

Where multiple control replicates were acquired, all sample replicates were normalized to each individual control replicate to account for total variability. Numerical processing: Microsoft Excel (Office Professional Plus 2019, v1808). Statistics: GraphPad Prism 8.4.2; specific tests are reported in the Figure Legends/Results. Figures were prepared in CorelDRAW Graphics Suite 2023 Update 2 (v24.5.0.731).

## 3. Results

### K562 cells, NK-92 cells, and primary NK cells tolerate STEM-CELLBANKER^®^ EX in culture

Since freezing with SCB requires a 100% concentration, omitting the washing step leads to residual SCB during co-incubation. To evaluate the feasibility of performing a no-wash killing assay with cryo-preserved cells in SCB, we first examined its cytotoxicity toward primary NK cells and the NK-92 cell line as effector cells, as well as K562 cells as the standard target cells during 24 h of culture. In addition to standard culture conditions, K562 target cells and primary NK cells were also tested at room temperature. For primary NK cells, assays were performed both in the presence of IL-2, reflecting standard cultivation conditions, and in its absence, as IL-2 is not included during the cytotoxicity assay itself. Since no major differences were observed, IL-2 supplementation was not evaluated for NK-92 cells.

At room temperature, K562 cells tolerated all tested SCB concentrations, including 100%, without substantial loss of viability (Figure 1A). In contrast, at 37 °C, mean relative viability decreased to approximately 86–88% with 25–75% SCB and further declined to 56% at 100% SCB. For primary NK cells, viability was markedly more sensitive to SCB exposure (Figure 1B). Both in the presence and absence of IL-2, viability declined to ∼80% at 25% SCB, to ∼25% at 50% SCB, and to <5% at 75–100% SCB. At room temperature, baseline viability was already reduced to roughly half of that observed under standard culture conditions (0% SCB). Addition of 25% SCB further halved viability, but interestingly, higher SCB concentrations appeared to be tolerated somewhat better relative to this diminished baseline. Because of the low tolerance of primary NK cells, lower SCB concentrations were examined. At 5–15% SCB, mean viability remained above 95%. Nevertheless, a statistically significant reduction compared to control was detected at all concentrations. The NK-92 cell line showed a similar overall trend to primary NK cells, but with somewhat greater resilience at intermediate concentrations (Figure 1C). At 25% SCB, viability dropped to ∼80%. At 50% and 75% SCB, viability was maintained at 72% and 55%, respectively. However, as with primary NK cells, virtually no viable cells remained at 100% SCB. When smaller increments were tested, viability remained above 95% at 5% and 10% SCB, but dropped more noticeably at 15% (84%) and 20% (82%), with statistically significant differences compared to the control for concentrations above 10%. Based on these results, 5% SCB was selected for subsequent assays. To operationalize this condition in a no-wash, in-tube format, we next defined the volume and dilution constraints for co-culture and acquisition.

**Figure 1.**
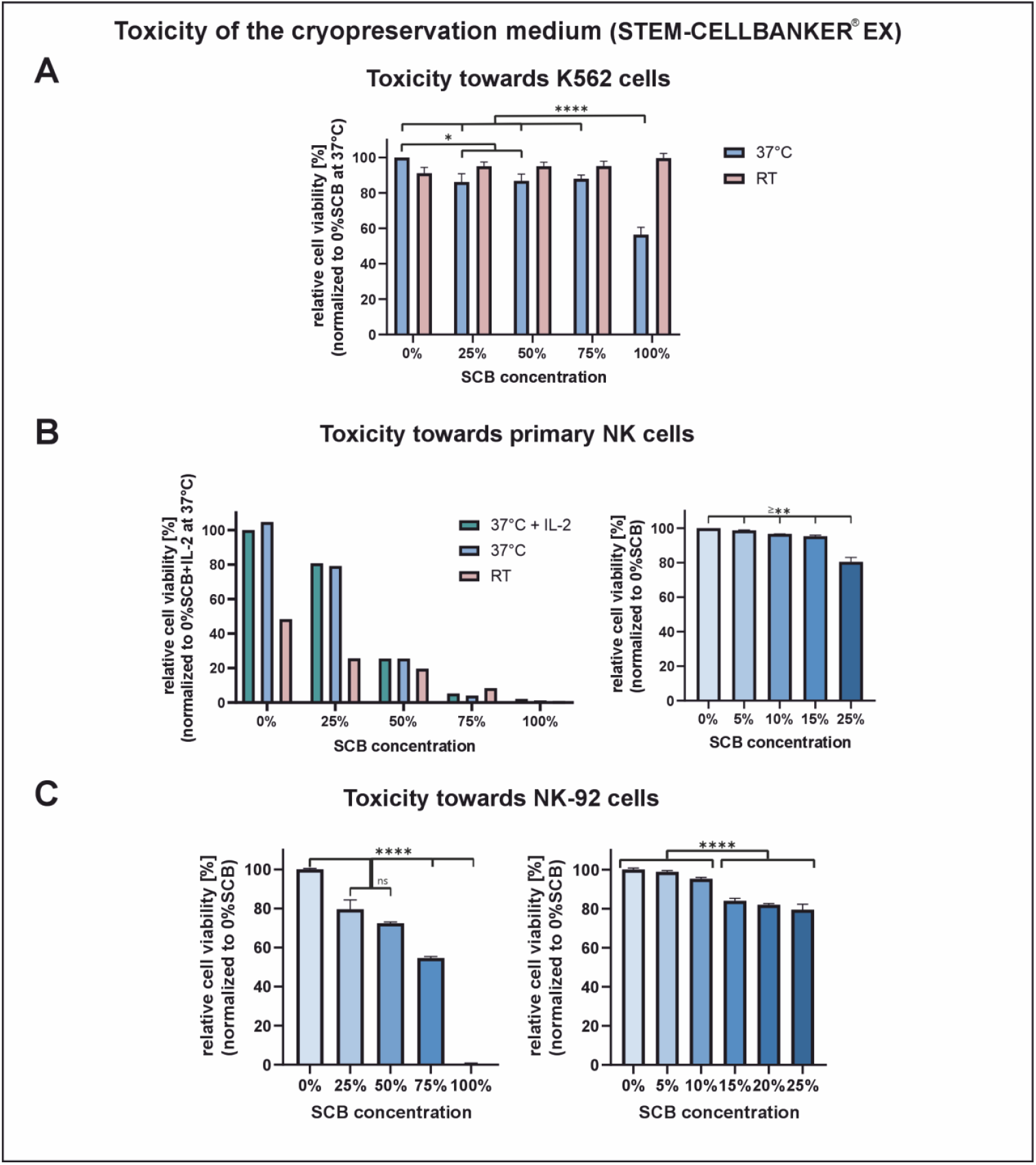
Toxicity of different concentrations of the cryopreservation medium STEM-CELLBANKER^®^ EX (SCB) towards K562 cells, primary NK cells, and NK-92 cells; cell viability was assessed via flow cytometry after 24 h incubation. **A:** Relative viability of K562 cells incubated with 0–100% SCB at 37°C and room temperature (RT) (n = 3 independent experiments, each 1 technical replicate); statistical analysis: two-way ANOVA with Tukey post hoc test within RT and 37°C groups. **B:** Relative viability of primary NK cells with 0–100% SCB at RT, 37°C, and 37°C + 500 U/mL IL-2 (left, n = 1 experiment, 1 technical replicate; no statistics performed) and with 0–25% SCB at 37°C (right, n = 2 donors/experiments, each 3 technical replicates); statistical analysis: two-way ANOVA with Tukey post hoc test. **C:** Relative viability of NK-92 cells with 0–100% SCB (left, n = 1 experiment, 3 technical replicates) and 0– 25% SCB (right, n = 1 experiment, 3 technical replicates) at 37°C; statistical analysis: one-way ANOVA with Tukey post hoc test. Bars represent mean ± SEM (ns = not significant; *p < 0.05; **p < 0.01; ****p < 0.0001).

### Viable K562 cells are well maintained after cryopreservation in STEM-CELLBANKER^®^ EX across densities

To achieve a final SCB concentration of <5%, a 20-fold dilution of the initial sample is required. Since our upper limit was defined as 800 µl during co-culture, this makes up to 40 µl of cryopreserved cells an appropriate starting volume. Since such a 20-fold dilution, combined with the loss of target cells due to NK cell-mediated killing, necessitates sufficiently high initial cell concentrations, we next examined how freezing density affects post-thaw viability. For this purpose, cells were cryopreserved at densities ranging from 1 × 10^6 to 5 × 10^6 cells (in increments of 1 × 10^6) in SCB, and their viability was assessed either directly in SCB or after washing.

Overall, viability was reduced by approximately 10% in samples that were not washed to remove SCB, a statistically significant effect (Figure 2A). Within both washed and unwashed groups, however, freezing density showed no consistent impact on viability (Figure 2B). Across all densities, washed cells exhibited a mean relative viability of 98% compared to pre-freezing levels, indicating that cryopreservation itself had only minor effects on viability. Mean viable recovery ranged from 73% to 85%, with averages of 80% in unwashed samples and 79% in washed samples; this difference was not statistically significant (Figure 2A). Mean total recovery ranged from 73% to 96%, with averages of 91% in unwashed samples and 81% in washed samples, which was statistically significant.

**Figure 2.**
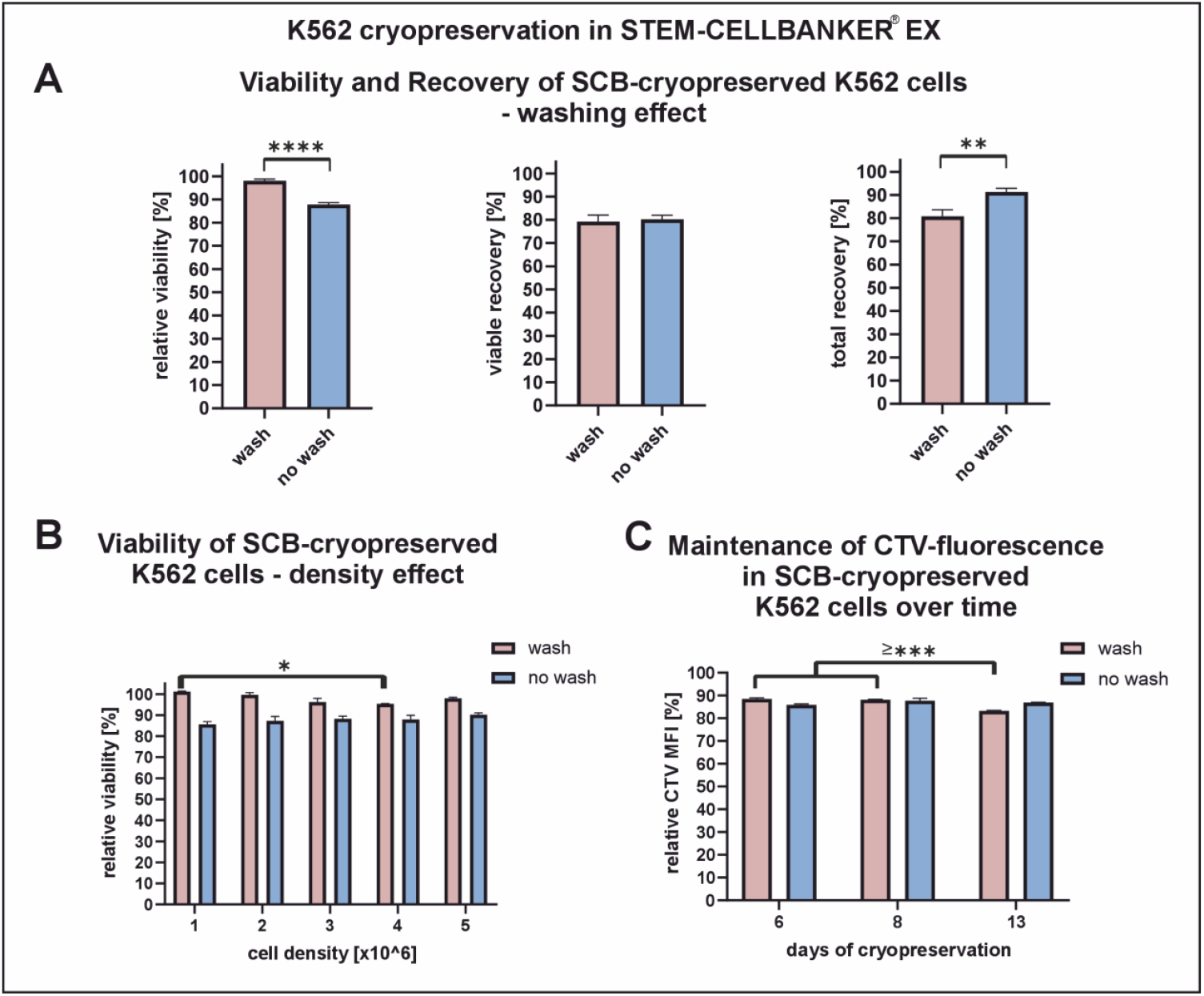
Effects of K562 cell cryopreservation in STEM-CELLBANKER^®^ EX (SCB) and post-thaw washing; all readouts normalized to pre-cryopreservation values. **(A**,**B)** K562 cells were cryopreserved at 1, 2, 3, 4, and 5 × 10^6 cells/mL in 1-mL cryotubes; after thawing, aliquots were either washed or not washed before flow-cytometric analysis. **A:** pooled relative viability (left), pooled viable recovery (middle), and pooled total recovery (right). **B:** relative viability shown separately for each freezing density (n = 1 experiment, 3 technical replicates); statistical analysis: two-way ANOVA with Tukey post hoc test. **C:** CellTrace™ Violet–labeled K562 cells were cryopreserved in 1-mL cryotubes and thawed at different time points; aliquots were washed or not washed prior to analysis (n = 1 experiment, 3 technical replicates); statistical analysis: two-way ANOVA with Tukey post hoc test. Bars represent mean ± SEM (*p < 0.05; **p < 0.01; ***p < 0.001; ****p < 0.0001).

### CTV labeling remains stable after cryopreservation with STEM-CELLBANKER^®^ EX

Because target identification is required for lysis quantification in mixed cultures, we next assessed the stability of target labeling after cryopreservation. We therefore validated whether CTV provides reliable target cell discrimination after cryopreservation. Fluorescence intensity was measured after 6, 8, and 13 days of storage. Relative mean fluorescence intensity ranged from 83% to 88% of pre-freeze levels (Figure 2C). In washed samples, fluorescence intensity on day 13 was significantly lower than on days 6 or 8, but no consistent temporal trend was observed. In unwashed samples, no significant differences were detected across timepoints. When comparing washed and unwashed conditions, significant differences were found at days 6 and 13, with higher fluorescence in unwashed cells, whereas no difference was observed on day 8. Overall, these data indicate that CTV labeling intensity decreases slightly after cryopreservation but remains stable across storage durations, enabling reliable discrimination of target cells in subsequent assays.

### Cytotoxic activity of NK-92 and primary NK cells is maintained in the presence of STEM-CELLBANKER^®^ EX

Having established both low cytotoxicity of SCB and stable target cell labeling after cryopreservation, we next assessed the impact of SCB on NK cell-mediated cytotoxicity. NK-92 cells and primary NK cells (cultured in the presence of IL-2) were tested against CTV-labeled K562 targets. K562 cells were cryopreserved in SCB, thawed, and either washed or left unwashed prior to co-culture. Effector and target cell densities were quantified by flow cytometry and adjusted to 5 × 10^6 viable cells/mL (targets in SCB, effectors in K562 medium). Cell suspensions were then combined at defined effector-to-target (E:T) ratios in K562 medium, resulting in a final SCB concentration of 5%, and co-cultured for 4 h. Following incubation, samples were cooled to 8 °C for 30 min before addition of PBS containing EDTA, FCS, PI, and CountBright™ beads. Flow cytometry was used to acquire the samples, and specific lysis was calculated either by (i) the proportion of viable versus dead targets (viability-based) or (ii) the absolute number of viable targets in a defined volume, as determined by bead normalization (bead-based) (Figure 3A).

**Figure 3.**
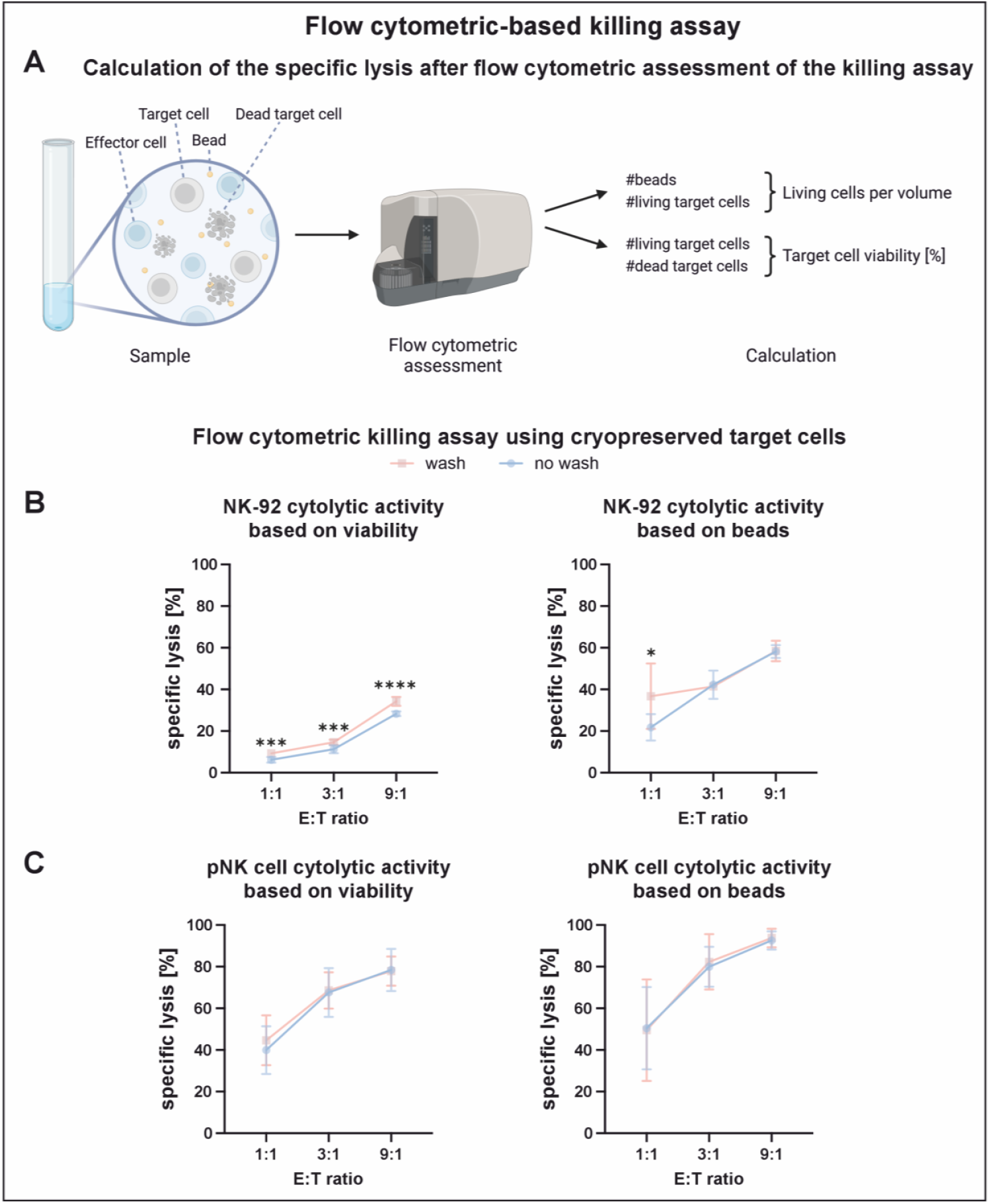
Analysis of specific lysis of washed versus non-washed target cells using bead-based absolute counting or viability-based calculation across effector-to-target (E:T) ratios. **A:** Schematic of the analysis workflow; after flow cytometric acquisition, specific lysis was calculated using either counting beads to derive viable targets per defined volume (bead-based) or the numbers of live and dead targets (viability-based). Created in BioRender. Kaltschmidt, C. (2025) https://BioRender.com/x49u069 **B:** Specific lysis in assays using primary NK (pNK) cells as effectors (left: viability-based; right: bead-based; n = 5 donors/experiments, each 2–3 technical replicates). **C:** Specific lysis in assays using NK-92 cells as effectors (left: viability-based; right: bead-based; n = 1 experiment, 3 technical replicates). Statistical analysis: two-way ANOVA with Sidak post hoc test. Bars represent mean ± SD (*p < 0.05; ***p < 0.001; ****p < 0.0001).

Two-way ANOVA confirmed that both the calculation method (p < 0.0001) and the effector-to-target (E:T) ratio (p < 0.0001) were the primary determinants of measured lysis. As expected, increasing effector cell numbers resulted in significantly higher lysis across both calculation approaches (Figure 3B and C). For NK-92 cells with washed targets, lysis values at E:T ratios of 1:1 and 1:3 were not significantly different, despite the overall trend. In primary NK cells, the increase from 1:1 to 1:3 was consistently significant, whereas the further step from 1:3 to 1:9 plateaued and did not reach significance in several donors (three in washed and one in unwashed conditions for bead-based analysis; two in washed and two in unwashed conditions for viability-based analysis). Bead-based calculations, which quantify the absolute number of remaining viable targets, consistently yielded higher lysis values than viability-based calculations, with the largest differences observed in NK-92 assays.

In contrast, the presence or absence of a washing step had only minor effects. In NK-92 assays, viability-based calculations showed small but reproducible differences of 3–6% across all E:T ratios (all p < 0.0001), whereas bead-based calculations detected a difference only at the 1:1 ratio (global effect p < 0.05) (Figure 3B). For primary NK cells, wash effects were inconsistent: significance was observed in only a single donor/ratio combination per calculation method, contributing just 0.1% to the overall variance (Figure 3C). Although the ANOVA identified a significant interaction term, suggesting donor-specific variation in wash effects, the contribution to overall variance was minimal. In summary, while both the calculation method and the E:T ratio had strong and consistent impacts on measured lysis, washing had only minimal and inconsistent effects. Furthermore, the presence of 5% SCB did not significantly influence NK-mediated cytotoxicity under either calculation approach (Figure 3B and C).

### Ready-to-use aliquot format: Micronic 1.1 mL tubes enable no-wash, in-tube acquisition

Having confirmed the feasibility of a killing assay using thawed target cells without washing steps, we next aimed to establish a format in which target cells could be cryopreserved as ready-to-use aliquots. To ensure compatibility with standard flow cytometry workflows, we sought a tube format that is cryopreservable, fits into conventional flow cytometry tubes, and can be stored in racks with a 96-well format. For this purpose, we selected 1.1 mL Micronic tubes featuring an external thread, a 96-well rack system, and a V-shaped bottom. Although these tubes fit seamlessly into standard flow cytometry tubes, the raised bottom caused interference with the sample pickup tube of the Beckman Coulter Gallios cytometer. Rather than modifying the autosampler setup, we deferred resolving this issue and temporarily transferred samples into alternative tubes for measurement.

### Thawing and volume optimization: direct warm-medium thaw equals water-bath; viability plateaus ≥80 µL

With the necessity of a 20-fold dilution, the volume of frozen target cell aliquots must be kept sufficiently small. As the diameter of cryopreservation tubes does not scale with volume, varying fill volumes alter the surface area-to-volume ratio, which can in turn affect freezing and thawing dynamics. To address this, we cryopreserved K562 cells in different volumes and assessed post-thaw viability and recovery. In addition, we compared two thawing procedures: conventional thawing in a water bath and direct thawing by addition of pre-warmed medium, which is more practical when handling multiple tubes in parallel. Overall viability was low in both conditions, but no significant difference was observed between the two thawing methods (Figure 4A). In both cases, higher fill volumes tended to yield improved viability, although the effect did not reach statistical significance. Based on these findings, we continued with the more convenient pre-warmed medium approach and extended the analysis by including larger volumes to further explore this trend.

**Figure 4.**
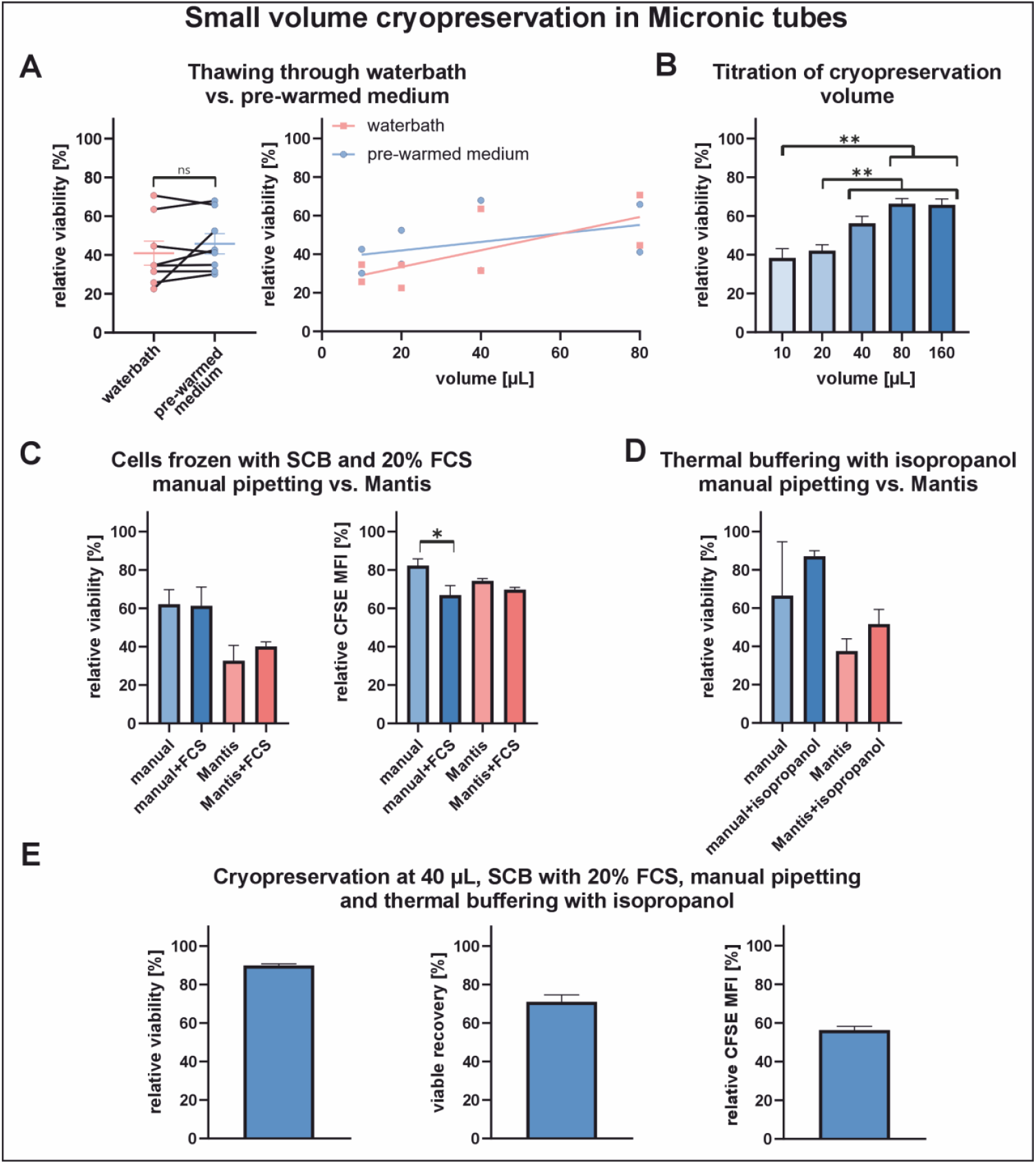
Optimization of small-volume cryopreservation in Micronic tubes; all readouts normalized to pre-cryopreservation values. **A:** Left, thawing by water bath versus direct addition of pre-warmed medium yielded comparable viability; right, post-thaw viability increased with cryo-volume (n = 1 experiment, 1 technical replicate, 2 time points); statistical analysis: paired t test (left) and simple linear regression (right). **B:** Post-thaw viability plateaued above ∼80 µL (n = 3 experiments, 1 technical replicate, 3 time points); statistical analysis: repeated-measures one-way ANOVA with Sidak post hoc test. **C:** Addition of 20% FCS to SCB produced a non-significant increase in mean post-thaw viability in cells dispensed by the automated system (left) but slightly reduced CFSE fluorescence retention (right) (n = 1 experiment, 5 technical replicates); statistical analysis: ordinary one-way ANOVA with Tukey post hoc test (both). **D:** Thermal buffering with isopropanol insulation during freezing resulted in a non-significant increase in post-thaw viability (n = 1 experiment, 2–5 technical replicates: 2 for manual, 3 for manual+isopropanol, 5 for both Mantis conditions); statistical analysis: ordinary one-way ANOVA with Sidak post hoc test. **E:** Relative viability (left), viable recovery (middle), and CFSE relative MFI (right) under the optimized small-volume cryo-preservation parameters. Bars represent mean ± SEM (ns = not significant; *p < 0.05; **p < 0.01).

A trend toward improved viability with increasing volumes was again observed; however, a plateau appeared to be reached at 80 µL (Figure 4B). One-way ANOVA confirmed no statistically significant differences between 40 µL, 80 µL, and 160 µL. We therefore selected 40 µL as the working volume. Accordingly, for subsequent analyses, we switched to a Cytek Northern Lights cytometer, which accommodates the selected tube format through software-controlled adjustment of the pickup tube depth. Since the available instrument configuration included only a 488 nm laser, target cell staining was changed from CTV to CFSE. In addition, the integrated volumetric measurement capability eliminated the need for counting beads. We evaluated the use of the Mantis^®^ Liquid Dispenser (Formulatrix), an automated system providing greater speed and accuracy compared with manual pipetting. In parallel, we tested whether supplementing SCB with 20% FCS could improve viability. In addition, this experiment served to validate the applicability of CFSE as an alternative to CTV. Relative viability of manually pipetted samples remained above 60% and was not affected by the addition of FCS (Figure 4C). In contrast, viability of samples dispensed with the Mantis was lower at 33%, but increased to a mean of 40% upon FCS supplementation. These differences, however, did not reach statistical significance. Relative MFI of CFSE in viable cells remained above 65% across all samples, with a slight but statistically significant reduction observed in manually pipetted samples upon FCS addition. In summary, FCS supplementation slightly reduced CFSE MFI in manually pipetted samples but showed a non-significant trend toward higher viability in Mantis-dispensed samples. 20% FCS was therefore included in the subsequent cryopreservation protocol.

To improve the relatively low viable recovery, we tested the placement of heat-sealed isopropanol-filled bags inside the Styrofoam box around the Micronic tube rack to serve as a thermal buffer, similar to commercial cryopreservation containers such as the Nalgene^®^ Mr. Frosty. The addition of isopropanol bags did not result in statistically significant differences, although both conditions showed a trend toward improved viability, leading to their inclusion in the cryopreservation protocol (Figure 4D). Dispensing with the Mantis again resulted in lower viabilities, albeit without statistical significance, and manual pipetting was therefore retained (Figure 4C and D). These optimizations were then combined into a consolidated protocol for pilot testing.

### Consolidated protocol performance: 90% relative viability and 71% viable recovery

With the cryopreservation procedure established—40 µl volume, manual pipetting, 20% FCS in SCB, and insulation with isopropanol-filled bags—we performed an additional experiment to assess relative viability and viable recovery under a streamlined, single-condition setup. Relative viability reached 90 ± 2% (mean ± SD) and viable recovery 71 ± 7% (mean ± SD) (Figure 4E). The relative CFSE MFI of viable cells was 56 ± 4% (mean ± SD); therefore, to maintain clear separation of targets and effectors, the CFSE staining concentration was increased in subsequent killing assays.

Based on a mean viable recovery of ∼70%, the targeted viable cell concentration of 5 × 10^6 cells/mL was adjusted to a cryopreservation density of 7.15 × 10^6 cells/mL. After thawing, mean relative viability was 74%, while absolute viability decreased from 86% before freezing to 63% after thawing (Figure 5A). During co-culture in the killing assay, relative viability showed a slight but statistically significant increase (p < 0.05).

**Figure 5.**
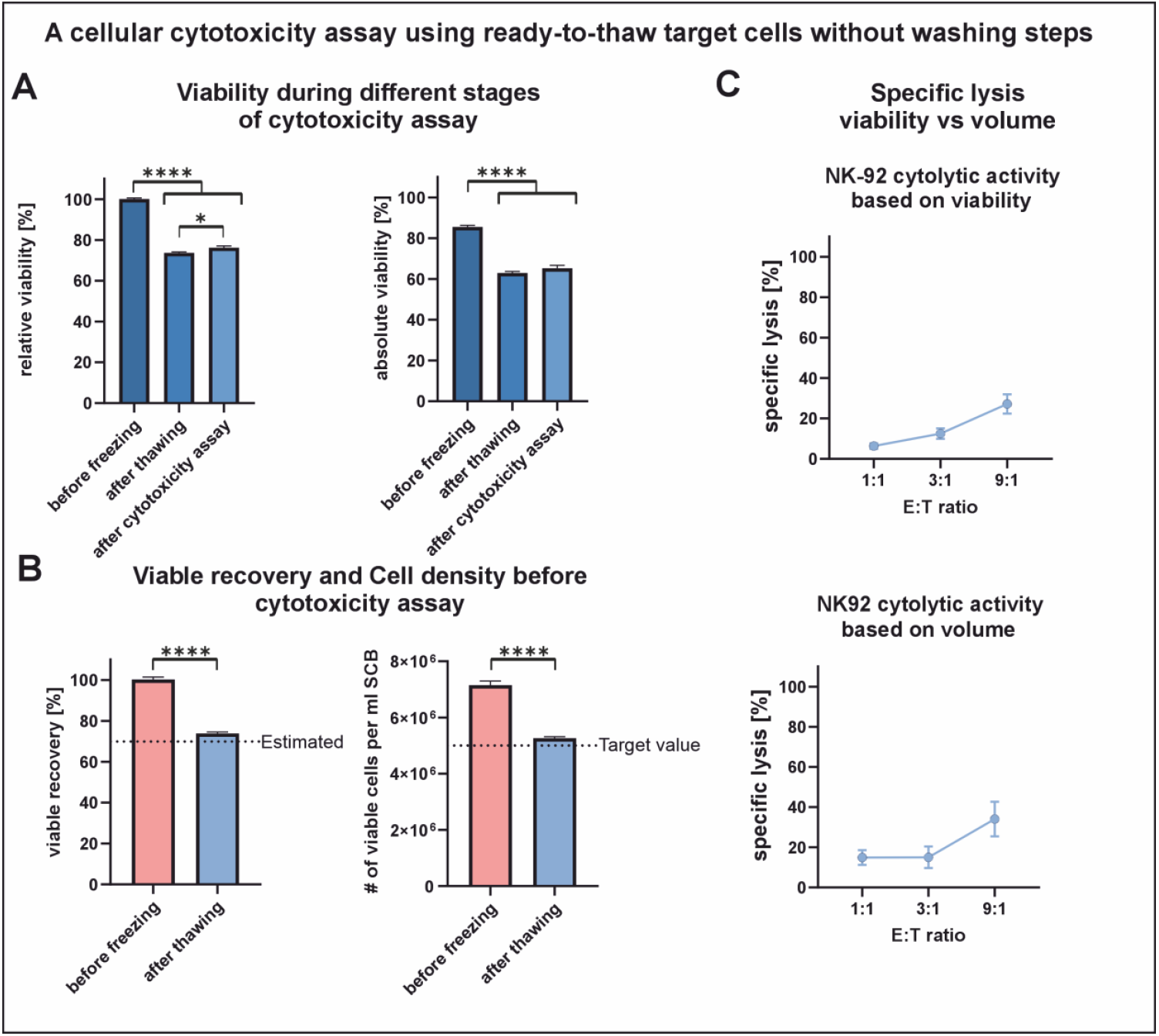
Cellular cytotoxicity assay using ready-to-thaw target cells without washing. CFSE-stained K562 targets were frozen in 40 µL aliquots of SCB with 20% FCS using isopropanol thermal buffering, thawed by direct addition of pre-warmed medium, and co-incubated with NK-92 cells for 4 h; EDTA and propidium iodide were then added for conjugate dissociation and dead-cell staining. **A:** Relative (left) and absolute (right) viability of target cells without NK-92 before cryopreservation, after thawing, and after the cytotoxicity assay (n = 1 experiment, 4–5 technical replicates: 5 before cryopreservation; 4 for both post-thaw conditions); statistical analysis: ordinary one-way ANOVA with Tukey post hoc test (both). **B:** Consistency of viable recovery (left) enables adjustment of pre-cryo-preservation target-cell density to achieve the desired post-thaw cell count (right) (n = 1 experiment, 4–5 technical replicates: 5 before cryopreservation; 4 after thawing); statistical analysis: unpaired t test (both). **C:** Specific lysis measured with NK-92 effectors by viability-based calculation (upper) and by volumetric absolute counts (lower) (n = 1 experiment, 4–5 technical replicates per E:T ratio: 4 for 0:1 and 9:1; 5 for 1:1 and 3:1). Bars represent mean ± SEM. (*p < 0.05; ****p < 0.0001).

### Assay-to-assay comparison: viability-based agreement; volumetric versus bead normalization differs at 1:3 and 1:9

The mean viable recovery of 74 ± 3% (mean ± SD) was within the expected range of 70 ± 7% (mean ± SD) (Figure 5B). As a result, the actual cell concentration at the start of the cytotoxicity assay was 5.3 × 10^6 cells/mL, slightly higher than the targeted value. At the 1:1 ratio, this resulted in a deviation of 5 ± 2% (mean ± SD) in the effective ratio, which diminished to ∼1% at the 1:3 and 1:9 ratios. When calculated from viability, specific lysis in the ready-to-thaw format was within the range observed in the prior no-wash assay (Figure 5 C, see Figure 3B). We then compared viable-count–based lysis between assays. At E:T ratios of 1:3 and 1:9, values differed significantly, indicating a discrepancy between bead-based (prior assay) and cytometer-based volumetric (ready-to-thaw) calculations, whereas viability-based lysis showed no difference.

## 4. Discussion

Cytotoxicity assays are essential tools for characterizing NK- and T cells in research, quality control, and translational applications [25,34,35]. As noted above, there is a need for a standardized, reproducible, and practical format that can be applied independently of freshly prepared target cells. Existing alternatives, such as enzymatic, fluorescence-based, or reporter cell line systems, address some limitations of classical methods but do not provide a standardized, ready-to-use solution [26]. To date, no “ready-to-thaw, no-wash” format with pre-labeled, cryopreserved target cells has been available to enable direct and robust application in cellular cytotoxicity assays.

Our results demonstrate that this concept is feasible: (i) at low residual concentrations, SCB is well tolerated by both effector and target cells and preserves viability (Figure 1); (ii) washing to remove SCB is not required and does not diminish assay performance (Figure 3); (iii) after cryopreservation, target-cell labeling remains sufficient for reliable discrimination (Figures 2 and 4); (iv) the choice of lysis calculation (bead/volumetric vs. viability gating) exerts a greater influence on reported cytotoxicity than other procedural variables (Figure 3); and (v) optimization of aliquot volume and supplementation established practical cryopreservation parameters that enable assay-ready 40-µL aliquots and allow no-wash assays to be run directly in the same tubes (Figures 4 and 5).

SCB is marketed as a Good Manufacturing Practice (GMP)-grade, xeno-free, chemically defined cryopreservation medium composed of inactive ingredients used in intravenously administered products and compatible with simple freezing workflows that do not require a controlled-rate freezer [32]. Based on this positioning, we hypothesized that low SCB concentrations would be well tolerated in cell culture. This was confirmed experimentally: K562 targets showed high tolerance, including at elevated SCB levels (Figure 1A), whereas primary NK and NK-92 cells were more sensitive; nevertheless, viability in both effector types remained ≥95% at ≤10% SCB (Figure 1B–C). We performed 24-h tolerability testing, whereas the cytotoxicity assay itself runs for 4 h. This 24-h tolerance profile supports extended-duration variants, such as overnight whole-blood NK cytotoxicity assays and long-duration T-cell killing assays that commonly quantify killing over durations of 4 h up to ≥24 h [26,36–42]. We therefore set 5% as the target residual SCB level for this format. This threshold preserved ≥95% viability across effectors (Figure 1B–C), maintained assay performance over 4 h (Figures 1, 3), and left dilution headroom for future variants.

In standard 1-mL cryovials, cryopreservation of K562 cells with SCB was well tolerated (Figure 2). Peer-reviewed studies report use of STEM-CELLBANKER formulations across multiple human cell types, supporting the general suitability of this medium family, although we did not identify publications explicitly using the EX variant [43–50]. Washing after thawing increased apparent viability but did not improve viable recovery, while reducing total recovery (living + dead) (Figure 2A). This may affect cytotoxicity assays that rely on viability percentages but not those based on the absolute loss of viable targets measured through beads or volumetric counting. Varying the freezing density between 1 × 10^6 and 5 × 10^6 cells/mL produced no detectable trend in post-thaw viability (Figure 2B); this range is within the SCB EX datasheet recommendation [31]. Because no density effect was observed within 1–5 × 10^6 cells/mL, higher concentrations appear feasible. In line with common practice (cell-type dependent), peripheral blood mononuclear cell (PBMC) protocols often freeze at 5–10 × 10^6 cells/mL, and some guides allow up to ∼5 × 10^7 cells/mL per 1-mL cryovial [51–54].

To achieve a ready-to-use, pre-aliquoted format, we set an upper working volume of ∼1 mL per tube; downstream additions (e.g., EDTA and a viability dye) further constrained the co-culture volume to ∼800 µL. Because low residual SCB is beneficial for viability, we titrated fill volumes and compared direct addition of pre-warmed medium with conventional water-bath thawing. As no difference was detected (Figure 4A), we adopted the direct-addition approach for improved operability. This is consistent with the general principle that thawing should be as rapid as possible to minimize solution effects in the critical temperature zone, which is particularly relevant for small volumes with low thermal capacity [53,55,56]. We subsequently identified 40 µL as a pragmatic fill volume at which doubling the volume did not improve viability significantly (Figure 4A–B). The 40-µL aliquot (i) maintains <5% residual SCB during co-culture, (ii) leaves sufficient space for reagents, and (iii) avoids the need for transfer into different tubes for measurement. Adding 20% of FCS produced a non-significant trend toward higher viability—consistent with the well described broad cytoprotective effects of serum supplements and their widespread use in cryopreservation [57–62]. Because heat is transmitted from the inside out, surface-area-to-volume ratio governs cooling dynamics in small aliquots. The principal challenge is uncontrolled supercooling, which depresses recovery and increases well-to-well variance; controlling ice nucleation markedly improves outcomes in small-volume or plate-based cryopreservation [63–66]. Although controlled-rate freezing is preferable when available [57,66,67], many laboratories rely on low-cost passive methods. We used a Styrofoam box at −70 to −80 °C and observed robust performance with 1-mL cryovials in SCB (Figure 2), whereas very small volumes did not reach similarly high viabilities (Figure 4). To stabilize cooling in our 96-rack Micronic tubes, we placed heat-sealed, isopropanol-filled bags around the rack inside the Styrofoam container, simulating the thermal buffering of devices such as Mr. Frosty (Nalgene); this produced a non-significant positive trend, plausibly by moderating cooling toward ∼1 °C/min [55,56]. While automated dispensing offers clear throughput advantages, use of the Mantis^®^ liquid dispenser led to reduced viability under our conditions. Given its common use for dispensing cell suspensions [68,69], this likely reflects device-specific shear/transient exposure or process timing that can be optimized; evaluation with alternative systems (e.g., a Hamilton Microlab-type pipetting robot) may mitigate these effects.

To discriminate targets from effectors without biasing the readout, we labeled the target population. Labeling effector cells can overestimate cytotoxicity because viable targets engaged in effector– target conjugates may be excluded during gating; labeling targets or adding a brief EDTA step at the end of culture mitigates this issue [25]. Consistently, flow-cytometric NK assays commonly pre-label targets to ensure unambiguous enumeration [35]. CTV labeling remained stable after cryopreservation in our hands (relative MFI 83–88% vs. pre-freeze across days 6–13), providing sufficient separation for reliable discrimination (Figure 2). This aligns with published data showing that CTV-labeled PBMC banks remain stable for up to 509 days of storage [70]. After switching to a 488-nm-only cytometer, we replaced CTV with CFSE; post-thaw brightness was lower (∼56% relative MFI), necessitating a higher staining concentration to maintain clear target–effector separation (Figures 4–5). Although cryopreservation can modestly affect signal, the observed MFI drop is also compatible with the known proliferation-independent intensity loss within the first 24 h after labeling, which is particularly pronounced for CFSE relative to CTV [71,72]. Thus, while our strategy yielded robust gating, alternative dye choices or a short post-labeling culture phase (optionally under division arrest) could further stabilize signal for CFSE-based labeling.

To quantify specific lysis in flow-cytometric cytotoxicity assays, two readouts can be used. The first is a viability-based measure, where specific lysis is calculated from the change in the fraction of viable versus dead targets between co-culture and control (i.e., specific lysis in % ≈ [(viable fraction in control - viable fraction in co-culture)/viable fraction in control] × 100). This readout is straightforward and performs well at lower E:T ratios, but it can underestimate lysis when killed targets are fragmented or lost from the target gate. Because target loss is a known issue, the second readout—absolute enumeration of remaining live targets—is generally preferred. Here, absolute counts are obtained either by adding counting beads (targets-per-bead or bead-normalized volume) or by using a cytometer with intrinsic volumetric metering that reports the analyzed volume directly (i.e., specific lysis in % ≈ 100-(viable fraction in co-culture/viable fraction in control) × 100) [73]. Because these approaches capture physical loss of targets in addition to overt death, they typically yield higher lysis values than viability gating under conditions of substantial killing; similar behavior is observed when direct enumeration methods are compared with release-based assays, which can underestimate specific lysis [17]. In our dataset, the calculation method and the E:T ratio were the principal determinants of reported lysis (two-way ANOVA), and absolute-count readouts consistently exceeded viability-based values (Figure 3). When we transitioned from bead normalization to a volumetric cytometer, small but systematic differences appeared at higher E:T ratios (3:1 and 9:1), underscoring that the counting strategy itself is a material source of variance and must be controlled when comparing datasets (Figures 3 and 5). Note that volumetric readings are not always considered exact and may need calibration, while bead-based single-platform methods introduce pipetting-related variance and require thorough mixing to prevent bead settling and time-dependent drift (reverse pipetting and vigorous mixing are standard recommendations) [73–78]. Overall, a no-wash workflow reduces operator effects and simplifies standardization across runs and laboratories—an approach aligned with single-platform absolute counting practices that minimize sample manipulation [79]. In primary NK co-cultures we observed a plateau in specific lysis between E:T = 3:1 and 9:1 (Figure 3C), consistent with saturating/kinetic limits within a fixed 4-h window (e.g., finite conjugation rates, serial-killing capacity, or target depletion in small volumes). Practically, extending incubation time often yields more dynamic range than further increasing E:T at short time scales [80–83].

Across both readouts in cytotoxicity assays with primary NK cells, omitting the post-thaw wash did not change the cytotoxicity outcome in a consistent direction (Figure 3C). In line with NK-92 cells being more susceptible to SCB (Figure 1), the viability-based readout showed small but statistically significant differences at every E:T ratio (Figure 3B). Mechanistically, adding a similar number of dead events to both samples can slightly inflate percent lysis in a viability-fraction calculation, leading to expectation of lower lysis levels in the washed condition. However, the net effect was very small; accordingly, the absolute-count readout (bead/volumetric) did not register differences beyond noise at 3:1 and 9:1. A donor-wash interaction reached significance (two-way ANOVA) but accounted for only a minimal fraction of variance, and—with no consistent pattern across donors or E:T ratios—likely reflects minor setup variance rather than a systematic biological effect. Overall, omitting the wash step has, at most, a minor impact on cytolytic function and does not alter overall effect trends. Practically, removing the wash reduces operator-dependent variability, eliminates tube transfers, and shortens turnaround, while preserving the biological signal and avoiding recovery loss from handling. It also harmonizes with our assay-ready 40-µL aliquots and in-tube acquisition, where residual SCB is kept ≤5% and labeling remains sufficient for discrimination (Figures 2, 4–5), facilitating more robust cross-run and cross-laboratory comparisons.

### Limitations and outlook

This study was scoped to K562 targets with NK-92 and primary NK effectors, a 4-h readout window, two target dyes (CTV/CFSE), and a small-volume, in-tube workflow based on SCB. We did not benchmark alternative cryomedia, evaluate active ice-nucleation control below 40 µL, or fully optimize automated dispensing; results were generated in a single laboratory and on a limited set of donors/instruments. Methodologically, minor differences between bead-based and volumetric enumeration at higher E:T ratios highlight that the chosen counting strategy can shape the reported lysis and should be kept constant when comparing datasets.

Conceptually, the format is transferable. Beyond SCB, other dimethyl sulfoxide (DMSO)-containing or DMSO-free cryomedia should be compatible provided the residual carryover is empirically titrated for effectors, targets, and dyes. Iterative tuning of the co-culture medium (e.g., protein supplementation, buffering/osmolarity, ionic strength) may reduce the dilution required post-thaw, enabling even smaller working volumes and potentially true 96-well implementations when combined with ice-nucleation control for sub-40 µL aliquots. On the analytical side, longer assay windows (overnight NK; ≥24 h T-cell killing) can extend dynamic range when primary cells exhibit E:T plateaus at short time scales. The workflow should also generalize in biological scope. Although demonstrated with cryo-preserved targets, cryopreserved effectors are feasible if conjugates are efficiently disrupted before acquisition (e.g., EDTA-containing buffers). With appropriate safeguards to prevent pre-freeze contact, “all-in-one” tubes containing both targets and effectors could be used to screen activating/inhibitory modulators or third components (e.g., bystander cells for tolerability and activating components like antibodies or bispecific antibodies). Variants using PBMCs or whole blood are straightforward, and additional readouts (e.g., degranulation) are compatible with the format; wash-free detection may require adapted reagents but does not conflict with the core workflow. Finally, adapting targets beyond K562—including adherent tumor cells or primary cancer stem-cell models—should be possible with format-specific handling (e.g., assay-ready suspensions, plate-based monolayers, or spheroids/organoids; using gentle dissociation where needed, such as with dispase or collagenase) and the same principles for labeling and absolute enumeration.

## 5. Conclusions

We establish a ready-to-thaw, no-wash flow-cytometric cytotoxicity assay that uses pre-labeled, cryopreserved target cells and can be run directly in-tube. At ≤5% residual SCB, both effectors and targets preserved viability, and post-thaw washing was unnecessary for assay performance. Target labeling (CTV/CFSE) remained sufficient after cryopreservation, and absolute enumeration (bead-normalized or volumetric) proved more robust than viability gating—making counting strategy, rather than washing, the dominant analytical determinant. Workflow optimization yielded assay-ready 40-µL aliquots compatible with direct acquisition, reducing handling steps, acquisition transfers, and operator variability. In primary NK assays, increasing E:T beyond 3:1 offered limited additional dynamic range within a 4 h window, supporting the use of fixed, moderate E:T together with longer incubations when deeper sensitivity is required. Collectively, these elements convert cytotoxicity testing from a labor-intensive, day-of-preparation procedure into a standardized, reproducible format suitable for research pipelines and translational quality control, and provide a practical template for broader adoption and future extensions.

## Author Contributions

Conceptualization, J.S., N.K., C.K. (Cornelius Knabbe), B.K. and C.K. (Christian Kaltschmidt); methodology, J.S. and N.K.; formal analysis, J.S., N.K., M.G., C.F., L.B., and J.B.; investigation, J.S., N.K., M.G., C.F., L.B., and J.B.; resources, C.K. (Cornelius Knabbe), B.K. and C.K. (Christian Kaltschmidt); data curation, J.S. and N.K.; writing—original draft preparation, J.S., N.K., M.G., C.F., L.B., and J.B.; writing—review and editing, J.S., N.K., M.G., C.F., L.B., J.B., C.K. (Cornelius Knabbe), B.K. and C.K. (Christian Kaltschmidt); visualization, J.S. and N.K.; supervision, C.K. (Cornelius Knabbe), B.K. and C.K. (Christian Kaltschmidt); project administration, J.S. and N.K.; funding acquisition, C.K. (Cornelius Knabbe), B.K. and C.K. (Christian Kaltschmidt) All authors have read and agreed to the published version of the manuscript.

## Institutional Review Board Statement

The study was conducted in accordance with the Declaration of Helsinki and approved by the Ethics Committee of the HDZ NRW, Department of Medicine, Ruhr University of Bochum (registry no. 2022-916, 7 March 2022).

## Acknowledgments

The authors thankfully acknowledge the Microscopy and Imaging Core Facility (Bielefeld University) for providing access to the Beckman Coulter Gallios™ flow cytometer. Furthermore, the authors thank the Forschungsverbund Biomedizin Bielefeld / OWL e.V. for providing access to the Cytek Northern Lights™ flow Cytometer donated by the Marlies und Herbert Repkow Stiftung (Welle 8, 33602 Bielefeld). During the preparation of this manuscript, the authors used OpenAI ChatGPT5 for the purposes of assisting with consolidating and editing the text (spell check, grammar check, harmonizing terminology, resolving internal inconsistencies, and standardizing reagent/vendor notation). The authors have reviewed and edited the output and take full responsibility for the content of this publication.

## Abbreviations

The following abbreviations are used in this manuscript:

ANOVA: Analysis of variance
BPxx: Band-pass filter (xx-nm center)
CAR: Chimeric antigen receptor
CAR-NK: Chimeric antigen receptor natural killer cells
CAR-T: Chimeric antigen receptor T cells
CFSE: Carboxyfluorescein succinimidyl ester
CTV: CellTrace™ Violet
CTFR: CellTrace™ Far Red
DAPI: 4′,6-diamidino-2-phenylindole
DCSP: Dichroic short-pass
DMSO: Dimethyl sulfoxide
E:T: Effector-to-target ratio
EDTA: Ethylenediaminetetraacetic acid
FBS: Fetal bovine serum
FCS: Fetal calf serum
FL1: Fluorescence channel 1
FL3: Fluorescence channel 3
FL6: Fluorescence channel 6
FL9: Fluorescence channel 9
FSC-A: Forward scatter area
FSC-H: Forward scatter height
GMP: Good Manufacturing Practice
IL-2: Interleukin-2
LDH: Lactate dehydrogenase
MEM-α: Minimum Essential Medium, alpha modification
MFI: Mean fluorescence intensity
NK: Natural killer
PBMC: Peripheral blood mononuclear cells
PBS: Phosphate-buffered saline
PBSE2%FCS: PBS + 2 mM EDTA + 2% FCS
PI: Propidium iodide
QC: Quality control
RPMI-1640: Roswell Park Memorial Institute 1640 medium
RT: Room temperature
SCB: STEM-CELLBANKER® EX
SD: Standard deviation
SEM: Standard error of the mean
SSC-A: Side scatter area

## References

1. Zhang, X.; Zhu, L.; Zhang, H.; Chen, S.; Xiao, Y. CAR-T Cell Therapy in Hematological Malignancies: Current Opportunities and Challenges. Front. Immunol. 2022, 13, 927153, doi:10.3389/fimmu.2022.927153.

2. Lambert, N.; El Moussaoui, M.; Baron, F.; Maquet, P.; Darcis, G. Virus-Specific T-Cell Therapy for Viral Infections of the Central Nervous System: A Review. Viruses 2023, 15, doi:10.3390/v15071510.

3. Fung, J.S.T.; Wright, R.C.; Bharaj, D.K.; Alghamdi, A.; Hesson, D.; Delisle, J.S.; Schweitzer, L.; Avery, R.K.; Belga, S. Virus-specific T-cell therapy for prophylaxis and treatment of cytomegalovirus infections after transplantation: a scoping review. Clin. Infect. Dis. 2025, doi:10.1093/cid/ciaf232.

4. Neller, M.A.; Ambalathingal, G.R.; Hamad, N.; Sasadeusz, J.; Pearson, R.; Holmes-Liew, C.-L.; Singhal, D.; Tunbridge, M.; Ng, W.Y.; Sharplin, K.; et al. Compassionate access to virus-specific T cells for adoptive immunotherapy over 15 years. Nat. Commun. 2024, 15, 10339, doi:10.1038/s41467-024-54595-2.

5. Page, A.; Chuvin, N.; Valladeau-Guilemond, J.; Depil, S. Development of NK cell-based cancer immunotherapies through receptor engineering. Cell. Mol. Immunol. 2024, 21, 315–331, doi:10.1038/s41423-024-01145-x.

6. Zhong, Y.; Liu, J. Emerging roles of CAR-NK cell therapies in tumor immunotherapy: current status and future directions. Cell Death Discov. 2024, 10, 318, doi:10.1038/s41420-024-02077-1.

7. Shi, Y.; Hao, D.; Qian, H.; Tao, Z. Natural killer cell-based cancer immunotherapy: from basics to clinical trials. Exp. Hematol. Oncol. 2024, 13, 101, doi:10.1186/s40164-024-00561-z.

8. Ahmad, A. CAR-T Cell Therapy. Int. J. Mol. Sci. 2020, 21, doi:10.3390/ijms21124303.

9. Bhaskar, S.T.; Dholaria, B.; Savani, B.N.; Sengsayadeth, S.; Oluwole, O. Overview of approved CAR-T products and utility in clinical practice. Clin. Hematol. Int. 2024, 6, 93–99, doi:10.46989/001c.124277.

10. Goyco Vera, D.; Waghela, H.; Nuh, M.; Pan, J.; Lulla, P. Approved CAR-T therapies have reproducible efficacy and safety in clinical practice. Hum. Vaccin. Immunother. 2024, 20, 2378543, doi:10.1080/21645515.2024.2378543.

11. Liu, E.; Marin, D.; Banerjee, P.; Macapinlac, H.A.; Thompson, P.; Basar, R.; Nassif Kerbauy, L.; Overman, B.; Thall, P.; Kaplan, M.; et al. Use of CAR-Transduced Natural Killer Cells in CD19-Positive Lymphoid Tumors. N. Engl. J. Med. 2020, 382, 545–553, doi:10.1056/NEJMoa1910607.

12. Shimasaki, N.; Jain, A.; Campana, D. NK cells for cancer immunotherapy. Nat. Rev. Drug Discov. 2020, 19, 200–218, doi:10.1038/s41573-019-0052-1.

13. Lazarski, C.A.; Hanley, P.J. Review of flow cytometry as a tool for cell and gene therapy. Cytotherapy 2024, 26, 103–112, doi:10.1016/j.jcyt.2023.10.005.

14. Marton, C.; Clémenceau, B.; Dachy, G.; Demerle, C.; Derenne, S.; Ferrand, C.; Giverne, C.; Latouche, J.-B.; Lemée, L.; Martinet, J.; et al. Harmonisation of quality control tests for academic production of CAR-T cells: a position paper from the WP-bioproduction of the UNITC consortium. Bone Marrow Transplant. 2025, 60, 1209–1217, doi:10.1038/s41409-025-02637-8.

15. Shao, L.; Zheng, Y.; Somerville, R.P.; Stroncek, D.F.; Jin, P. New insights on potency assays from recent advances and discoveries in CAR T-cell therapy. Front. Immunol. 2025, 16, 1597888, doi:10.3389/fimmu.2025.1597888.

16. Brunner, K.T.; Mauel, J.; Cerottini, J.C.; Chapuis, B. Quantitative assay of the lytic action of immune lymphoid cells on 51-Cr-labelled allogeneic target cells in vitro; inhibition by isoantibody and by drugs. Immunology 1968, 14, 181–196.

17. Somanchi, S.S.; McCulley, K.J.; Somanchi, A.; Chan, L.L.; Lee, D.A. A Novel Method for Assessment of Natural Killer Cell Cytotoxicity Using Image Cytometry. PLoS One 2015, 10, e0141074, doi:10.1371/journal.pone.0141074.

18. Karimi, M.A.; Lee, E.; Bachmann, M.H.; Salicioni, A.M.; Behrens, E.M.; Kambayashi, T.; Baldwin, C.L. Measuring cytotoxicity by bioluminescence imaging outperforms the standard chromium-51 release assay. PLoS One 2014, 9, e89357, doi:10.1371/journal.pone.0089357.

19. Yang, L.; Shen, M.; Xu, L.J.; Yang, X.; Tsai, Y.; Keng, P.C.; Chen, Y.; Lee, S.O. Enhancing NK cell-mediated cytotoxicity to cisplatin-resistant lung cancer cells via MEK/Erk signaling inhibition. Sci. Rep. 2017, 7, 7958, doi:10.1038/s41598-017-08483-z.

20. Zaritskaya, L.; Shurin, M.R.; Sayers, T.J.; Malyguine, A.M. New flow cytometric assays for monitoring cell-mediated cytotoxicity. Expert Rev. Vaccines 2010, 9, 601–616, doi:10.1586/erv.10.49.

21. Phan, M.-T.; Chun, S.; Kim, S.-H.; Ali, A.K.; Lee, S.-H.; Kim, S.; Kim, S.-H.; Cho, D. Natural killer cell subsets and receptor expression in peripheral blood mononuclear cells of a healthy Korean population: Reference range, influence of age and sex, and correlation between NK cell receptors and cytotoxicity. Hum. Immunol. 2017, 78, 103–112, doi:10.1016/j.humimm.2016.11.006.

22. Lorenzo-Herrero, S.; Sordo-Bahamonde, C.; Gonzalez, S.; López-Soto, A. CD107a Degranulation Assay to Evaluate Immune Cell Antitumor Activity. Methods Mol. Biol. 2019, 1884, 119–130, doi:10.1007/978-1-4939-8885-3_7.

23. Li, Y.; Yu, M.; Yin, J.; Yan, H.; Wang, X. Enhanced Calcium Signal Induces NK Cell Degranulation but Inhibits Its Cytotoxic Activity. J. Immunol. 2022, 208, 347–357, doi:10.4049/jimmunol.2001141.

24. Lion, E.; Smits, E.L.J.M.; Berneman, Z.N.; van Tendeloo, V.F.I. Quantification of IFN-gamma produced by human purified NK cells following tumor cell stimulation: comparison of three IFN-gamma assays. J. Immunol. Methods 2009, 350, 89–96, doi:10.1016/j.jim.2009.08.014.

25. Wu, X.; Zhang, Y.; Li, Y.; Schmidt-Wolf, I.G.H. Improvements in Flow Cytometry-Based Cytotoxicity Assay. Cytometry A 2021, 99, 680–688, doi:10.1002/cyto.a.24242.

26. Kim, J.; Phan, M.-T.T.; Kweon, S.; Yu, H.; Park, J.; Kim, K.-H.; Hwang, I.; Han, S.; Kwon, M.-J.; Cho, D. A Flow Cytometry-Based Whole Blood Natural Killer Cell Cytotoxicity Assay Using Overnight Cytokine Activation. Front. Immunol. 2020, 11, 1851, doi:10.3389/fimmu.2020.01851.

27. LIVE/DEAD™ Cell-Mediated Cytotoxicity Kit, for animal cells 1 Kit | Invitrogen™. Available online: https://www.thermofisher.com/order/catalog/product/L7010 (accessed on 28 September 2025).

28. 7-AAD/CFSE Cell-Mediated Cytotoxicity Assay Kit | Cell Signaling Technology. Available online: https://www.cellsignal.com/products/cellular-assay-kits/7-aad-cfse-cell-mediated-cytotoxicity-assay-kit/72782?srsltid=AfmBOoo0TUH1jaIBPNPb_Q-k9diBBswoqLgpep5w6eyTTLTTyJvZoi (accessed on 28 September 2025).

29. Tocris Bioscience. FlowX Human NK Cell Killing Flow Cytometry Kit. Available online: https://www.rndsystems.com/products/flowx-human-nk-cell-killing-flow-cytometry-kit_fmc032 (accessed on 28 September 2025).

30. Target Cell Killing Bioassays. Available online: https://www.promega.de/en/products/reporter-bioas-says/target-cell-killing-bioassays/ (accessed on 28 September 2025).

31. STEM-CELLBANKER® EX Cryopreservation Solution | Amsbio. Datasheet. Available online: https://resources.amsbio.com/Datasheets/11936.pdf (accessed on 27 September 2025).

32. STEM-CELLBANKER® EX Cryopreservation Solution | Amsbio. Available online: https://www.ams-bio.com/stem-cell-cryopreservation-ex/ (accessed on 27 September 2025).

33. Kusch, N.; Storm, J.; Macioszek, A.; Knabbe, C.; Kaltschmidt, B.; Kaltschmidt, C. Donor Variability and Seeding Density Shape NK-Cell Proliferation and Surface Receptor Expression: Insights from an Integrated Phenotypic and Genetic Analysis. Cells 2025, 14, doi:10.3390/cells14161252.

34. Galdina, V.; Puga Yung, G.L.; Seebach, J.D. Cytotoxic Responses Mediated by NK Cells and Cytotoxic T Lymphocytes in Xenotransplantation. Transpl. Int. 2025, 38, 13867, doi:10.3389/ti.2025.13867.

35. Kandarian, F.; Sunga, G.M.; Arango-Saenz, D.; Rossetti, M. A Flow Cytometry-Based Cytotoxicity Assay for the Assessment of Human NK Cell Activity. J. Vis. Exp. 2017, doi:10.3791/56191.

36. Haeseleer, F.; Eichholz, K.; Tareen, S.U.; Iwamoto, N.; Roederer, M.; Kirchhoff, F.; Park, H.; Okoye, A.A.; Corey, L. Real-Time Killing Assays to Assess the Potency of a New Anti-Simian Immunodeficiency Virus Chimeric Antigen Receptor T Cell. AIDS Res. Hum. Retroviruses 2020, 36, 998–1009, doi:10.1089/AID.2020.0163.

37. Santos, J.; Ogando, J.; Lacalle, R.A.; Mañes, S. A flow cytometry-based method to screen for modulators of tumor-specific T cell cytotoxicity. Methods Enzymol. 2020, 631, 467–482, doi:10.1016/bs.mie.2019.02.040.

38. Wang, Z.; Wang, J.; Zhao, Y.; Jin, J.; Si, W.; Chen, L.; Zhang, M.; Zhou, Y.; Mao, S.; Zheng, C.; et al. 3D live imaging and phenotyping of CAR-T cell mediated-cytotoxicity using high-throughput Bessel oblique plane microscopy. Nat. Commun. 2024, 15, 6677, doi:10.1038/s41467-024-51039-9.

39. Chanda, M.K.; Shudde, C.E.; Piper, T.L.; Zheng, Y.; Courtney, A.H. Combined analysis of T cell activation and T cell-mediated cytotoxicity by imaging cytometry. J. Immunol. Methods 2022, 506, 113290, doi:10.1016/j.jim.2022.113290.

40. Jedema, I.; van der Werff, N.M.; Barge, R.M.Y.; Willemze, R.; Falkenburg, J.H.F. New CFSE-based assay to determine susceptibility to lysis by cytotoxic T cells of leukemic precursor cells within a heterogeneous target cell population. Blood 2004, 103, 2677–2682, doi:10.1182/blood-2003-06-2070.

41. Noto, A.; Ngauv, P.; Trautmann, L. Cell-based flow cytometry assay to measure cytotoxic activity. J. Vis. Exp. 2013, e51105, doi:10.3791/51105.

42. Jin, Q.; Jiang, L.; Chen, Q.; Li, X.; Xu, Y.; Sun, X.; Zhao, Z.; Wei, L. Rapid flow cytometry-based assay for the evaluation of γδ T cell-mediated cytotoxicity. Mol. Med. Rep. 2017, 17, 3555–3562, doi:10.3892/mmr.2017.8281.

43. Park, J.J.; Lee, O.-H.; Park, J.-E.; Cho, J. Comparison of Cryopreservation Media for Mesenchymal Stem Cell Spheroids. Biopreserv. Biobank. 2024, 22, 486–496, doi:10.1089/bio.2023.0057.

44. Leitner, D.; Ramamoorthy, M.; Dejosez, M.; Zwaka, T.P. Immature mDA neurons ameliorate motor deficits in a 6-OHDA Parkinson’s disease mouse model and are functional after cryopreservation. Stem Cell Res. 2019, 41, 101617, doi:10.1016/j.scr.2019.101617.

45. Baek, S.-K.; Cho, Y.-S.; Kim, I.-S.; Jeon, S.-B.; Moon, D.-K.; Hwangbo, C.; Choi, J.-W.; Kim, T.-S.; Lee, J.-H. A Rho-Associated Coiled-Coil Containing Kinase Inhibitor, Y-27632, Improves Viability of Dissociated Single Cells, Efficiency of Colony Formation, and Cryopreservation in Porcine Pluripotent Stem Cells. Cell. Reprogram. 2019, 21, 37–50, doi:10.1089/cell.2018.0020.

46. Yamazaki, T.; Enosawa, S.; Tokiwa, T. Effect of cryopreservation on the appearance and liver function of hepatocyte-like cells in cultures of cirrhotic liver of biliary atresia. In Vitro Cell. Dev. Biol. Anim. 2018, 54, 401–405, doi:10.1007/s11626-018-0260-8.

47. Kusakisako, K.; Masatani, T.; Yada, Y.; Talactac, M.R.; Hernandez, E.P.; Maeda, H.; Mochizuki, M.; Tanaka, T. Improvement of the cryopreservation method for the Babesia gibsoni parasite by using commercial freezing media. Parasitol. Int. 2016, 65, 532–535, doi:10.1016/j.parint.2016.02.012.

48. Shimazu, T.; Mori, Y.; Takahashi, A.; Tsunoda, H.; Tojo, A.; Nagamura-Inoue, T. Serum- and xeno-free cryopreservation of human umbilical cord tissue as mesenchymal stromal cell source. Cytotherapy 2015, 17, 593–600, doi:10.1016/j.jcyt.2015.03.604.

49. Al-Saqi, S.H.; Saliem, M.; Quezada, H.C.; Ekblad, Å.; Jonasson, A.F.; Hovatta, O.; Götherström, C. Defined serum- and xeno-free cryopreservation of mesenchymal stem cells. Cell Tissue Bank. 2015, 16, 181–193, doi:10.1007/s10561-014-9463-8.

50. Saliem, M.; Holm, F.; Tengzelius, R.B.; Jorns, C.; Nilsson, L.-M.; Ericzon, B.-G.; Ellis, E.; Hovatta, O. Improved cryopreservation of human hepatocytes using a new xeno free cryoprotectant solution. World J. Hepatol. 2012, 4, 176–183, doi:10.4254/wjh.v4.i5.176.

51. Protocol for Cryopreserving PBMCs | STEMCELL Technologies. Available online: https://www.stem-cell.com/how-to-cryopreserve-pbmcs.html (accessed on 27 September 2025).

52. Higdon, L.E.; Lee, K.; Tang, Q.; Maltzman, J.S. Virtual Global Transplant Laboratory Standard Operating Procedures for Blood Collection, PBMC Isolation, and Storage. Transplant. Direct 2016, 2, e101, doi:10.1097/TXD.0000000000000613.

53. Browne, D.J.; Miller, C.M.; Doolan, D.L. Technical pitfalls when collecting, cryopreserving, thawing, and stimulating human T-cells. Front. Immunol. 2024, 15, 1382192, doi:10.3389/fimmu.2024.1382192.

54. Paige Etter; Melissa Austin. PBMC_SOP.

55. Whaley, D.; Damyar, K.; Witek, R.P.; Mendoza, A.; Alexander, M.; Lakey, J.R. Cryopreservation: An Overview of Principles and Cell-Specific Considerations. Cell Transplant. 2021, 30, 963689721999617, doi:10.1177/0963689721999617.

56. Baboo, J.; Kilbride, P.; Delahaye, M.; Milne, S.; Fonseca, F.; Blanco, M.; Meneghel, J.; Nancekievill, A.; Gaddum, N.; Morris, G.J. The Impact of Varying Cooling and Thawing Rates on the Quality of Cryo-preserved Human Peripheral Blood T Cells. Sci. Rep. 2019, 9, 3417, doi:10.1038/s41598-019-39957-x.

57. Uhrig, M.; Ezquer, F.; Ezquer, M. Improving Cell Recovery: Freezing and Thawing Optimization of Induced Pluripotent Stem Cells. Cells 2022, 11, doi:10.3390/cells11050799.

58. Zalomova, L.V.; Fesenko, E.E., JR. FBS-based cryoprotective compositions for effective cryopreservation of gut microbiota and key intestinal microorganisms. BMC Res. Notes 2024, 17, 168, doi:10.1186/s13104-024-06836-2.

59. Nazarpour, R.; Zabihi, E.; Alijanpour, E.; Abedian, Z.; Mehdizadeh, H.; Rahimi, F. Optimization of Human Peripheral Blood Mononuclear Cells (PBMCs) Cryopreservation. Int. J. Mol. Cell. Med. 2012, 1, 88–93.

60. Juhl, M.; Christensen, J.P.; Pedersen, A.E.; Kastrup, J.; Ekblond, A. Cryopreservation of peripheral blood mononuclear cells for use in proliferation assays: First step towards potency assays. J. Immunol. Methods 2021, 488, 112897, doi:10.1016/j.jim.2020.112897.

61. Mohamed, H.M.; Sundar, P.; Ridwan, N.A.A.; Cheong, A.J.; Mohamad Salleh, N.A.; Sulaiman, N.; Mh Busra, F.; Maarof, M. Optimisation of cryopreservation conditions, including storage duration and revival methods, for the viability of human primary cells. BMC Mol. Cell Biol. 2024, 25, 20, doi:10.1186/s12860-024-00516-6.

62. Erol, O.D.; Pervin, B.; Seker, M.E.; Aerts-Kaya, F. Effects of storage media, supplements and cryopreservation methods on quality of stem cells. World J. Stem Cells 2021, 13, 1197–1214, doi:10.4252/wjsc.v13.i9.1197.

63. Daily, M.I.; Whale, T.F.; Kilbride, P.; Lamb, S.; John Morris, G.; Picton, H.M.; Murray, B.J. A highly active mineral-based ice nucleating agent supports in situ cell cryopreservation in a high throughput format. J. R. Soc. Interface 2023, 20, 20220682, doi:10.1098/rsif.2022.0682.

64. Murray, K.A.; Kinney, N.L.H.; Griffiths, C.A.; Hasan, M.; Gibson, M.I.; Whale, T.F. Pollen derived macromolecules serve as a new class of ice-nucleating cryoprotectants. Sci. Rep. 2022, 12, 12295, doi:10.1038/s41598-022-15545-4.

65. Meneghel, J.; Kilbride, P.; Morris, G.J. Cryopreservation as a Key Element in the Successful Delivery of Cell-Based Therapies-A Review. Front. Med. (Lausanne) 2020, 7, 592242, doi:10.3389/fmed.2020.592242.

66. Daily, M.I.; Whale, T.F.; Partanen, R.; Harrison, A.D.; Kilbride, P.; Lamb, S.; Morris, G.J.; Picton, H.M.; Murray, B.J. Cryopreservation of primary cultures of mammalian somatic cells in 96-well plates benefits from control of ice nucleation. Cryobiology 2020, 93, 62–69, doi:10.1016/j.cryobiol.2020.02.008.

67. Wuchter, P. The EBMT Handbook: Hematopoietic Cell Transplantation and Cellular Therapies: Processing, Cryopreserving, and Controlling the Quality of HSC, 8th; Cham (CH), 2024, ISBN 9783031440793.

68. Hirokawa, Y.; Clarke, J.; Palmieri, M.; Tan, T.; Mouradov, D.; Li, S.; Lin, C.; Li, F.; Luo, H.; Wu, K.; et al. Low-viscosity matrix suspension culture enables scalable analysis of patient-derived organoids and tumoroids from the large intestine. Commun. Biol. 2021, 4, 1067, doi:10.1038/s42003-021-02607-y.

69. Zukas, K.; Cayford, J.; Serneo, F.; Atteberry, B.; Retter, A.; Eccleston, M.; Kelly, T.K. Rapid high-throughput method for investigating physiological regulation of neustrophil extracellular trap formation. J. Thromb. Haemost. 2024, 22, 2543–2554, doi:10.1016/j.jtha.2024.05.028.

70. Nicotra, T.; Desnos, A.; Halimi, J.; Antonot, H.; Reppel, L.; Belmas, T.; Freton, A.; Stranieri, F.; Mebarki, M.; Larghero, J.; et al. Mesenchymal stem/stromal cell quality control: validation of mixed lymphocyte reaction assay using flow cytometry according to ICH Q2(R1). Stem Cell Res. Ther. 2020, 11, 426, doi:10.1186/s13287-020-01947-6.

71. Tario, J.D. JR,; Conway, A.N.; Muirhead, K.A.; Wallace, P.K. Monitoring Cell Proliferation by Dye Dilution: Considerations for Probe Selection. Methods Mol. Biol. 2018, 1678, 249–299, doi:10.1007/978-1-4939-7346-0_12.

72. Wallace, P.K.; Tario, J.D.; Fisher, J.L.; Wallace, S.S.; Ernstoff, M.S.; Muirhead, K.A. Tracking antigen-driven responses by flow cytometry: monitoring proliferation by dye dilution. Cytometry A 2008, 73, 1019–1034, doi:10.1002/cyto.a.20619.

73. Saraiva, L.; Wang, L.; Kammel, M.; Kummrow, A.; Atkinson, E.; Lee, J.Y.; Yalcinkaya, B.; Akgöz, M.; Höckner, J.; Ruf, A.; et al. Comparison of Volumetric and Bead-Based Counting of CD34 Cells by Single-Platform Flow Cytometry. Cytometry B Clin. Cytom. 2019, 96, 508–513, doi:10.1002/cyto.b.21773.

74. DeRose, P.C.; Benkstein, K.D.; Elsheikh, E.B.; Gaigalas, A.K.; Lehman, S.E.; Ripple, D.C.; Tian, L.; Vreeland, W.N.; Welch, E.J.; York, A.W.; et al. Number Concentration Measurements of Polystyrene Submicrometer Particles. Nanomaterials (Basel) 2022, 12, doi:10.3390/nano12183118.

75. CountBright Absolute Counting Beads | Molecular Probes. Product information. Available online: https://tools.thermofisher.com/content/sfs/manuals/mp36950.pdf (accessed on 28 September 2025).

76. BD Biosciences. BD Trucount™ Controls | BD Biosciences. Product information. Available online: https://www.bdbiosciences.com/content/dam/bdb/product_assets/product_pdf/flowcytometryproduct/pdf_0/23-3537.pdf (accessed on 28 September 2025).

77. Walker, C.; Barnett, D. Flow rate calibration for absolute cell counting rationale and design. Curr. Protoc. Cytom. 2006, Chapter 6, Unit6.24, doi:10.1002/0471142956.cy0624s36.

78. Storie, I.; Sawle, A.; Goodfellow, K.; Whitby, L.; Granger, V.; Reilly, J.T.; Barnett, D. Flow rate calibration I: a novel approach for performing absolute cell counts. Cytometry B Clin. Cytom. 2003, 55, 1–7, doi:10.1002/cyto.b.10051.

79. Petriz, J.; Bradford, J.A.; Ward, M.D. No lyse no wash flow cytometry for maximizing minimal sample preparation. Methods 2018, 134-135, 149–163, doi:10.1016/j.ymeth.2017.12.012.

80. Li, H.-K.; Hsiao, C.-W.; Yang, S.-H.; Yang, H.-P.; Wu, T.-S.; Lee, C.-Y.; Lin, Y.-L.; Pan, J.; Cheng, Z.-F.; Lai, Y.-D.; et al. A Novel off-the-Shelf Trastuzumab-Armed NK Cell Therapy (ACE1702) Using Antibody-Cell-Conjugation Technology. Cancers (Basel) 2021, 13, doi:10.3390/cancers13112724.

81. Bhat, R.; Watzl, C. Serial killing of tumor cells by human natural killer cells--enhancement by therapeutic antibodies. PLoS One 2007, 2, e326, doi:10.1371/journal.pone.0000326.

82. Subedi, N.; van Eyndhoven, L.C.; Hokke, A.M.; Houben, L.; van Turnhout, M.C.; Bouten, C.V.C.; Eyer, K.; Tel, J. An automated real-time microfluidic platform to probe single NK cell heterogeneity and cytotoxicity on-chip. Sci. Rep. 2021, 11, 17084, doi:10.1038/s41598-021-96609-9.

83. Khazen, R.; Müller, S.; Lafouresse, F.; Valitutti, S.; Cussat-Blanc, S. Sequential adjustment of cytotoxic T lymphocyte densities improves efficacy in controlling tumor growth. Sci. Rep. 2019, 9, 12308, doi:10.1038/s41598-019-48711-2.

